## Supplementary material for "SARS-CoV-2 Nsp14 mediates the effects of viral infection on the host cell transcriptome": Table 15

**Table 15: list of the plasmids for expressing SARS-CoV-2 proteins used for the screening in Figure 1.**

| Addgene ID | Plasmid Name |
|---|---|
| 141391 | pLVX-EF1alpha-SARS-CoV-2-N-2xStrep-IRES-Puro |
| 141385 | pLVX-EF1alpha-SARS-CoV-2-E-2xStrep-IRES-Puro |
| 141386 | pLVX-EF1alpha-SARS-CoV-2-M-2xStrep-IRES-Puro |
| 141383 | pLVX-EF1alpha-SARS-CoV-2-orf3a-2xStrep-IRES-Puro |
| 141394 | pLVX-EF1alpha-SARS-CoV-2-orf10-2xStrep-IRES-Puro |
| 141379 | pLVX-EF1alpha-SARS-CoV-2-nsp13-2xStrep-IRES-Puro |
| 141395 | pLVX-EF1alpha-eGFP-2xStrep-IRES-Puro |
| 141373 | pLVX-EF1alpha-SARS-CoV-2-nsp7-2xStrep-IRES-Puro |
| 141369 | pLVX-EF1alpha-SARS-CoV-2-nsp4-2xStrep-IRES-Puro |
| 141393 | pLXV-EF1alpha-2xStrep-SARS-CoV-2-orf9c-IRES-Puro |
| 141376 | pLVX-EF1alpha-SARS-CoV-2-nsp10-2xStrep-IRES-Puro |
| 141378 | pLVX-EF1alpha-SARS-CoV-2-nsp12-2xStrep-IRES-Puro |
| 141387 | pLVX-EF1alpha-SARS-CoV-2-orf6-2xStrep-IRES-Puro |
| 141390 | pLVX-EF1alpha-SARS-CoV-2-orf8-2xStrep-IRES-Puro |
| 141368 | pLVX-EF1alpha-SARS-CoV-2-nsp2-2xStrep-IRES-Puro |
| 141381 | pLVX-EF1alpha-SARS-CoV-2-nsp15-2xStrep-IRES-Puro |
| 141377 | pLVX-EF1alpha-SARS-CoV-2-nsp11-2xStrep-IRES-Puro |
| 141370 | pLVX-EF1alpha-SARS-CoV-2-nsp5-2xStrep-IRES-Puro |
| 141374 | pLVX-EF1alpha-SARS-CoV-2-nsp8-2xStrep-IRES-Puro |
| 141389 | pLXV-EF1alpha-2xStrep-SARS-CoV-2-orf7b-IRES-Puro |
| 141367 | pLVX-EF1alpha-SARS-CoV-2-nsp1-2xStrep-IRES-Puro |
| 141384 | pLXV-EF1alpha-2xStrep-SARS-CoV-2-orf3b-IRES-Puro |
| 141371 | pLVX-EF1alpha-SARS-CoV-2-nsp5-C145A-2xStrep-IRES-Puro |
| 141388 | pLVX-EF1alpha-SARS-CoV-2-orf7a-2xStrep-IRES-Puro |
| 141375 | pLVX-EF1alpha-SARS-CoV-2-nsp9-2xStrep-IRES-Puro |
| 141380 | pLXV-EF1alpha-2xStrep-SARS-CoV-2-nsp14-IRES-Puro |
| 141392 | pLVX-EF1alpha-SARS-CoV-2-orf9b-2xStrep-IRES-Puro |
