## Supplementary material for "SARS-CoV-2 Nsp14 mediates the effects of viral infection on the host cell transcriptome": Table 16

**Table 16. List of the primers used for qPCRs**

| TARGET | FORWARD PRIMER (5'-3') | REVERSE PRIMER (5'-3') |
|---|---|---|
| GAPDH | TCACCAGGGCTGCTTTTAAC | GACAAGCTTCCCGTTCTCAG |
| TBP | AGTGACCCAGGGTGCCAT | TGAATAGGCTGTGGGGTCAG |
| SH2D2A | CCTGTCCTACGGAAGAGCTG | CCTTGCCCAATCACAGAGTT |
| FGF-18 | GGCAAGGAGACGGAATTCTA | ACACACACTCCTTGCTGGTG |
| CXCL8 | AGCTCTGTGTGAAGGTGCAG | TGGGGTGGAAAGGTTTGGAG |
| COL13A | TGCCTCTAACCTTGGATTGG | TCTGGCATTTTCAAAAAGTGAA |
| CDC42-circRNA | TGTTCTGCACTTACACAGGTCA | TTTTACCAACAGCACCATCG |
| VRK1- circRNA | GGCAAATTGGACCTCAGTGT | CTTCCAGCTTGAGCTGCTTT |
| MARCHF7-circRNA | ACAAATGAACCAAGCACACG | GGAGCTGGAAGGTTGAACAG |
| CDK1-circRNA | AAATGGAAACCAGGAAGCCTA | GGGTATGGTAGATCCGAGAGC |
| U6 | GCTTCGGCAGCACATATACTAAAAT | CGCTTCACGAATTTGCGTGTCAT |
| TBP pre-mRNA | TGCTGGTTATGACAGCTCGAT | CATGTTTCCGGCGATTGCAT |
| 18S rRNA | GCTTAATTTGACTCAACACGGGA | AGCTATCAATCTGTCAATCCTGTC |
| PAXIP1 (exon-exon) | GGAGGTCAAGTATTACGCGGT | TGAGGCTAGTGCATTGTAGGAA |
| PAXIP1 (exon-intron) | GCCCATTTTAATTTGTGTGGA | TGAGGCTAGTGCATTGTAGGAA |
| ZNF507 (exon-exon) | AACGAGGCTCGCGGAAGCAG | CCAACATGGCAACACTGCTA |
| ZNF507 (exon-intron) | CATGCCTGTAATTATCCTGTAACC | CCAACATGGCAACACTGCTA |
| SETD1A (exon-exon) | ACTTCCGCTGGTGGCCTA | GAACAAACACGGATTTGCATT |
| SETD1A (exon-intron) | ACTTCCGCTGGTGGCCTA | CTACCCTCACCGCTCTCTGG |
